# Single cell sequencing during the entire life cycle reveals cell type diversity in *Oikopleura dioica*, and pools of genes expressed in the house-producing epithelium

**DOI:** 10.64898/2026.03.31.715263

**Authors:** Anthony Leon, Simon Henriet, David Lagman, Sara Barberan Martin, Astrid Canal, Gaelle Alleon, Cyprille Lenfant, Anne Aasjord, Daniel Chourrout

## Abstract

In tunicates, larvaceans represent a fascinating case of evolution, where the chordate body plan has been maintained despite a rapidly evolving genome characterized by strong In contrast to other tunicates, larvaceans keep the chordate body plan during their entire life. They have acquired a highly specialized epithelium in charge of producing the “house”, a complex extracellular apparatus used for filter feeding in the plankton. To what extent the house and this epithelium represent true molecular innovations withing chordates is a question for which thorough transcriptomics can bring novel insights.

We conducted a developmental profiling of gene expression at the single-cell level in the larvacean *Oikopleura dioica*. We provide detailed descriptions of cellular transcriptomes associated with the house-synthesizing organ, which permits to define the molecular specifics of epithelial cell territories. We followed their emergence during development, and we identified genes that represent key candidate molecules for regulating the morphogenesis of the house-producing organ. Dynamic changes in gene expression and cell identities during major developmental transitions of the lifecycle illustrate that our dataset effectively allows access to the diversity of *O. dioica*’s cell types in embryos and in adults. The resources presented here constitute critical assets to investigate larvacean biology and evolution for mechanistic and comparative goals.

## Introduction

Single-cell RNA sequencing (scRNA-seq) has already made a significant contribution by revealing a large number of cell types based on a specific molecular signature, and by predicting the gene pools involved in their development and function. It also allows us to advance our understanding of organismal evolution, when large phylogenetic distances complicate the recognition of tissue and organ homologies [1, 2]. Anatomical diversity arises not only from modifications of homologous organs but also from the emergence of specific structures, which can be considered as lineage-specific innovations when their structural or molecular homology is not apparent or is only minimally so [3].

Within the chordates, how tunicates evolved remains enigmatic. Their phylogenetic position between anatomically more complex groups suggests a secondary simplification [4] that may have conserved most of the ancestral developmental mechanisms [5]. For ascidians, remarkable models for the observation of chordate embryos, scRNA seq has already allowed the identification of many cell types [1] which complemented large *in situ hybridization* (ISH) screens [6]. The comparison of later life stages is complicated by a drastic metamorphosis that virtually erases the chordate body plan. Among tunicates, larvaceans are distinguished by the maintenance of this plan until the adult stage. Their ecological importance as one of the most abundant organisms in plankton [7], and the very short life cycle of some larvacean species that facilitates genetic analysis, have motivated numerous studies in recent times. However, larvacean gene expression is known for only a minority of developmental genes. Genome analyses have also shown a divergence of their gene content vs other chordates [8], with occasional gene losses [9] and frequent gene duplications [10]. Gene sequences also evolved rapidly which sometimes makes it difficult to recognize gene orthologs.

Despite maintaining the chordate body plan, larvaceans have also innovated in several ways, and the most apparent one is the construction of the “house”, a complex extracellular apparatus used for concentrating and filtering food particles. The larvacean house is composed of cellulose (which all tunicates can synthesize) and several dozen highly glycosylated proteins called oikosins [11], whose sequences show no tangible homology with genes of other organisms. The components of the house are produced and secreted by a highly differentiated oikoplastic epithelium, divided into several cellular territories well known to anatomists [12]. The oikoplastic epithelium covers almost the entire trunk of the animal. Some studies, including experimental ones, have shown the involvement of homeodomain protein transcription factors in its development [13, 14]. There is little doubt that elucidating the evolutionary origin of the oikoplastic epithelium and house synthesis will greatly benefit from more thorough transcriptomics and proteomics.

This study reports a scRNA-seq protocol performed at multiple stages from blastula to adult in the model species *Oikopleura dioica*. It allows, during development and later life, the identification of the molecular signature of the main organs and cell types; and the diverse territories of the oikoplastic epithelium. Examining these signatures opens the possibility of studying the molecular mechanisms at work in larvaceans and comparing them with those of other chordates, in a context of particularly rapid evolution.

## Materials and Methods

### Sample acquisition

Founder specimens of *Oikopleura dioica* were primarily collected from fjords near Bergen, Norway (60°34′N, 5°02′E and 60°15′N, 5°20′E). Populations were maintained in the laboratory at 13–15°C for several generations, following the conditions of Bouquet *et al.* [15]. Embryonic and larval samples were produced by *in vitro* fertilization of spawned eggs as described previously [16]. Fertilized eggs were washed extensively to remove female debris and excess sperm, and placed at in filtered artificial seawater (FASW) at 15°C. We monitored developmental progression at regular intervals until 16 hours post fertilization (16 hpf). Batches that included more than 10% abnormal morphologies were discarded. Each targeted developmental stage (2hpf, 2.5hpf, 3hpf, 3.5hpf, 4hpf, 6hpf, 8hpf, 11hpf, 16hpf) is represented by pooling at least 20 distinct broods. Adult samples were collected from the laboratory culture 24h before the targeted developmental stage at 5, 6 or 7 days post-fertilization (dpf), and starved overnight in facility seawater.

### Preparation of cell suspensions

Pooled individuals were transferred to a glass salier and washed extensively with Calcium-free artificial seawater (CASW) at 15°C to remove debris. Larval samples (from 8 to 16hpf) were passed on 40 µm strainer. Adult samples were anaesthetized by dropwise addition of 10% MS222-CASW and washed with CASW immediately prior to dissociation to remove houses. Cell dissociation was conducted in glass saliers pre-rinsed with 1% BSA-CASW, by adding to the 0.2 ml sample pellet a 2 ml digestion cocktail pre-warmed at 37°C (2-6phf samples: 0.5% Pronase, 5 mM EDTA, CASW; 8-6hpf samples: 0.25% Pronase, 1.25% Trypsin, 10mg.ml^-1^ Trichoderma Cellulase, CASW). Samples were digested for 5 minutes (2-6hpf samples) or 10 minutes (8-6hpf and adult samples) at 25°C, and 1 ml of the reaction volume was removed. We produced a cell suspension from digested samples by pipetting the remaining solution in and out a 1 ml plastic tip. Samples were pipetted until complete tissue disruption (usually 5 to 10 minutes) and reactions were stopped by adding 3 ml of ice-cold quenching solution (0.5%BSA, 2mg.ml^-1^ Glycine, CASW). After an additional 1-minute time of pipetting the sample, complete dissociation of cell aggregates was confirmed under microscope. Cell suspensions were passed on a 20 µm strainer and collected in a 50 ml Falcon tube pre-rinsed with 1% BSA-CASW. Strainers were washed with Magnesium-free CASW (1mM EDTA, CASW) supplemented with 0.5% BSA to maximize cell recovery. Cells were collected after 300g centrifugation for 15 minutes at 25°C and gently resuspended in 1 ml 0.2% BSA-MCASW. For 8-16hpf samples, we performed an additional centrifugation and wash step with 0.2% BSA-MCASW. For adult samples, we resuspended cells in PBS supplemented with 0.7 M D-Mannitol. We counted cells and checked viability using Acridine orange and Propidium iodide stains in LUNA-FL dual fluorescence cell counter. Cell mortality was under 20% for embryonic and larval samples, and under 30% for adult ones. Except for adult samples, all cell suspensions were subsequently processed for fixation.

### Acetic acid-Methanol (ACME) fixation

We implemented a minor modification to the ACME procedure of Garcia-Castro *et al.* [17] during fixation and cryopreservation of *O. dioica* cell suspensions. To prevent osmotic shock, we substituted the original 1x PBS solution with a high-salt buffer (HPBS; 0.5 M NaCl, 50 µM KCl, 0.25M Na_2_SO_4_, 0.32 M NaHCO_3_, 1mM KH2PO4, 3 mM Na_2_HPO_4_, pH 7.4). Cryopreserved cells stored at -80°C were resuspended in 1%BSA-HPBS to a concentration of 500 cells.µl^-1^ before barcoding.

### Gene expression profiling with bulk RNA-seq

Unfertilized eggs were collected immediately after spawning, and after a wash with AFSW to remove maternal cell debris. Samples corresponding to developing animals were produced by *in vitro* fertilization of spawned eggs as described above, and left to develop in FASW at 15°C before collecting successive time points. For all samples, the material was sedimented in a 1.5 ml tube before removing seawater and snap-freezing in liquid N_2_. Frozen pellets were resuspended in lysis buffer and total RNA was extracted with the NucleoSpin RNA XS kit (Macherey-Nagel), without applying carrier RNA. We checked RNA quality with a Bioanalyzer (Agilent) instrument prior to preparation of barcoded Illumina libraries using the NEBNext Single Cell/Low Input RNA Library Prep Kit (New England Biolabs). Samples were pooled, and we sequenced two pools with six libraries each on a Miseq v3 flowcell using 2x300 bp pair-end sequencing. After adapter removal, reads were mapped against the *O. dioica* genome assembly ASM20953v1 (Genbank GCA_000209535.1) using HISAT2 [18].

### Single-cell RNA-seq

In the case of fixed samples, we loaded ∼10^4^ fixed cells per well on a Chromium Next GEM chip (10X Genomics) following manufacturer’s recommendations for the Chromium controller. In the case of live samples, we loaded ∼2.10^4^ cells per well on a Chromium GEM-X chip (10X Genomics) following manufacturer’s recommendations for the Chromium-X controller.

We assessed the quality of Illumina cDNA libraries with Qubit (ThermoFisher) and Bioanalyzer instruments, prior to 2x100 bp pair-end sequencing on a Novaseq SP flowcell at the Norwegian Sequencing Centre (Oslo, Norway). The reads were mapped against the *O. dioica* genome assembly ASM20953v1 (Genbank GCA_000209535.1) and processed with the 10x Genomics Cellranger 9.0.1 pipeline for alignment, barcode and UMI counting. Raw feature matrices were processed with Cellbender 0.3.2 [19]. Filtering and clustering were performed with the R-package Seurat v5, with lower- and higher-limit cutoff of 2.10^2^ and 4.10^3^ RNA count per cell. After filtering, 160.424 cells were retained, of which 133.287 were classified as singlets with DoubletFinder. We normalized the data using Seurat’s LogNormalize approach with a scale factor of 10.000, followed by identification of 2.000 variable features using the vst method. Data were scaled and subjected to principal component analysis (PCA). Batch correction was performed using the Harmony package prior to downstream clustering. Nearest neighbor graphs were constructed using the first 40 principal components, and clustering was performed at a resolution of 1.4. UMAP was used for visualization of both unintegrated and Harmony-corrected embeddings.

### Whole mount in situ hybridization (ISH) techniques

Animal samples were fixed overnight in 4% paraformaldehyde (PFA) at 4°C. For chromogenic detection of transcripts, we carried out digoxygenin-labelled (DIG) probe synthesis and hybridization as described by Søviknes *et al.* [20]. Antisense probe designs were checked with BLAST against the *O. dioica* genome assembly for non-specific targets. We used AP-anti DIG conjugates and BCIP/NBT for chromogenic reactions conducted between 3h to 16h at 25°C. Mounted samples were observed with a Zeiss Axio Scope A1 upright microscope. For detecting transcripts with Hybridization Chain Reaction (HCR), we used a python script [21] for selecting optimal hybridization targets based on BLAST results against the *O. dioica* genome assembly.

Amplifiers and buffers were purchased from Molecular Instruments (Los Angeles, CA), and we followed the manufacturer’s recommendations to carry out hybridizations. Mounted samples were observed with an Olympus FV3000 confocal microscope.

## Results

### Single-cell transcriptomes during development

To gain a comprehensive overview of gene expression during the *O. dioica* lifecycle, we established single-cell transcriptomes for twelve developmental stages; corresponding to early embryos before gastrulation (2, 2.5 and 3hpf), embryos between gastrulation and hatching (3.5 and 4hpf), hatched larvae (6 and 8hpf), larvae post tailshift (11-12hpf), juveniles (16hpf), and maturing adults (5, 6 and 7dpf). Our collection represents 133.287 single-cell transcriptomes which, after unsupervised clustering, are resolved into 60 robust cell clusters (Fig 1A). Gene ontology (GO) enrichment analysis of all clusters shows that the proportion of mitochondrial transcripts varied modestly among cell populations, and their expression did not indicate cellular stress. We conclude that the dissociation procedure had only negligible impact on single-cell transcriptomes, and that our dataset should reflect gene expression in live samples.

**Figure 1:**
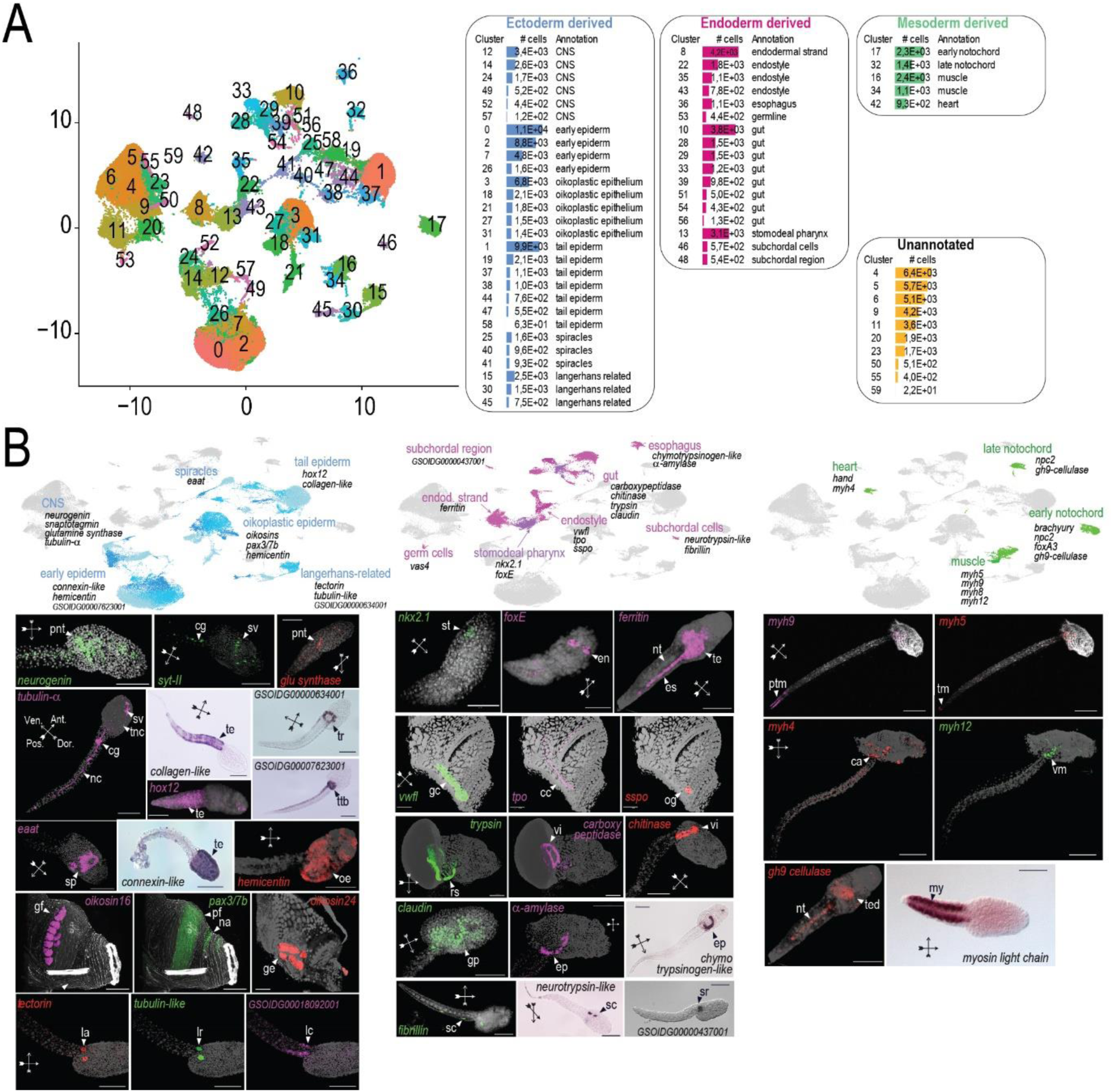
General annotation of *O. dioica* single-cell transcriptomes. **A)** Left panel, representation of all sequenced transcriptomes in low dimensional space. Numbers indicate cell clusters obtained from unsupervised clustering. Right panel, size, annotation and embryonic origin of cell clusters. **B)** Cluster annotation with detection of gene expression *in situ*. For each cluster examined, we produced antisense probes directed against transcripts of top-expressed candidate genes. Cluster identity was inferred from gene expression patterns observed in tissues of either developing embryos or adults: ca, cardiac cells; cc, corridor cells; cg, caudal ganglion; en, endostyle; ep, esophagus; es, endodermal strand; gc, glandular cells; ge, giant Eisen; gf, giant Fol; gp, gastric primordium; la, Langerhans receptor-associated cell; lc, lateral cells; lr, Langerhans mechanosensory receptor; my, tail muscle; na, Nasse cells; nc, nerve chord; nt, notochord; oe, oikoplastic epithelium; pnt, proneural territory; ptm, pre-tip tail muscle; rs, right stomach; sc, subchordal cell; sp, spiracles; sr, subchordal origin region; st, stomodeum; sv, sensory vesicle; te, tail epithelium; ted, trunk endoderm; tm, tail tip muscle; tnc, trunk nerve chord; tr, tail ring; ttb, tail-trunk boundary; vi, vertical intestine; vm, vesicular mesoderm.

Combined with the examination of homologous gene expressions, systematic annotation using *in situ* hybridization to target transcripts confirmed the identity of 49 clusters from the merged dataset. The annotation includes clusters representative of the major structures of *O. dioica* - for instance the oikoplastic epithelium, organs of the digestive tracts, and organs of the tail (Fig 1B). We also annotated clusters that correspond either to small organs composed by only few cells in larvae and adults, such as the Langerhans sensory receptor [22] and subchordal cells [23]; or to structures that are only transient during development, such as the endodermal strand [24]. These observations suggest that the collection successfully captures the diversity of cell types, even though fragile cells and cells present in fibrous structures that are difficult to dissociate like the adult tail, could be under-represented. For subsequent analyses aimed at monitoring developmental progression, stage-specific datasets were clustered individually.

### Functional annotation of house-producing cell territories

Prior to gonad growth, the oikoplastic epithelium represents a conspicuous organ of post-larval *O. dioica* specimens. It consists of a monolayer organized in distinct territories corresponding to highly differentiated cells (Fig 2A), some of which could attain much larger sizes through genome endoreduplications [25]. The oikoplastic epithelium is a hallmark of larvaceans, based on the ancient acquisition of the cellulose synthase gene *cesA2* [26] and on the *oikosin* genes that encode transmembrane and secreted glycoproteins [27–29] (Fig 2B). Single-cell transcriptomes of adult *O. dioica* provided us with a unique opportunity to examine further the molecular characteristics of the oikoplastic epithelium. We performed dimensionality reduction on adult cells expressing *cesA2*, to produce a variety of clusters that correspond to candidate oikoplastic cells (Fig 2E). Based on characteristic expression patterns of oikosins that were reported at the mRNA or protein level in distinct oikoplastic territories, we could allocate most clusters of this dataset to described cell types.

**Figure 2:**
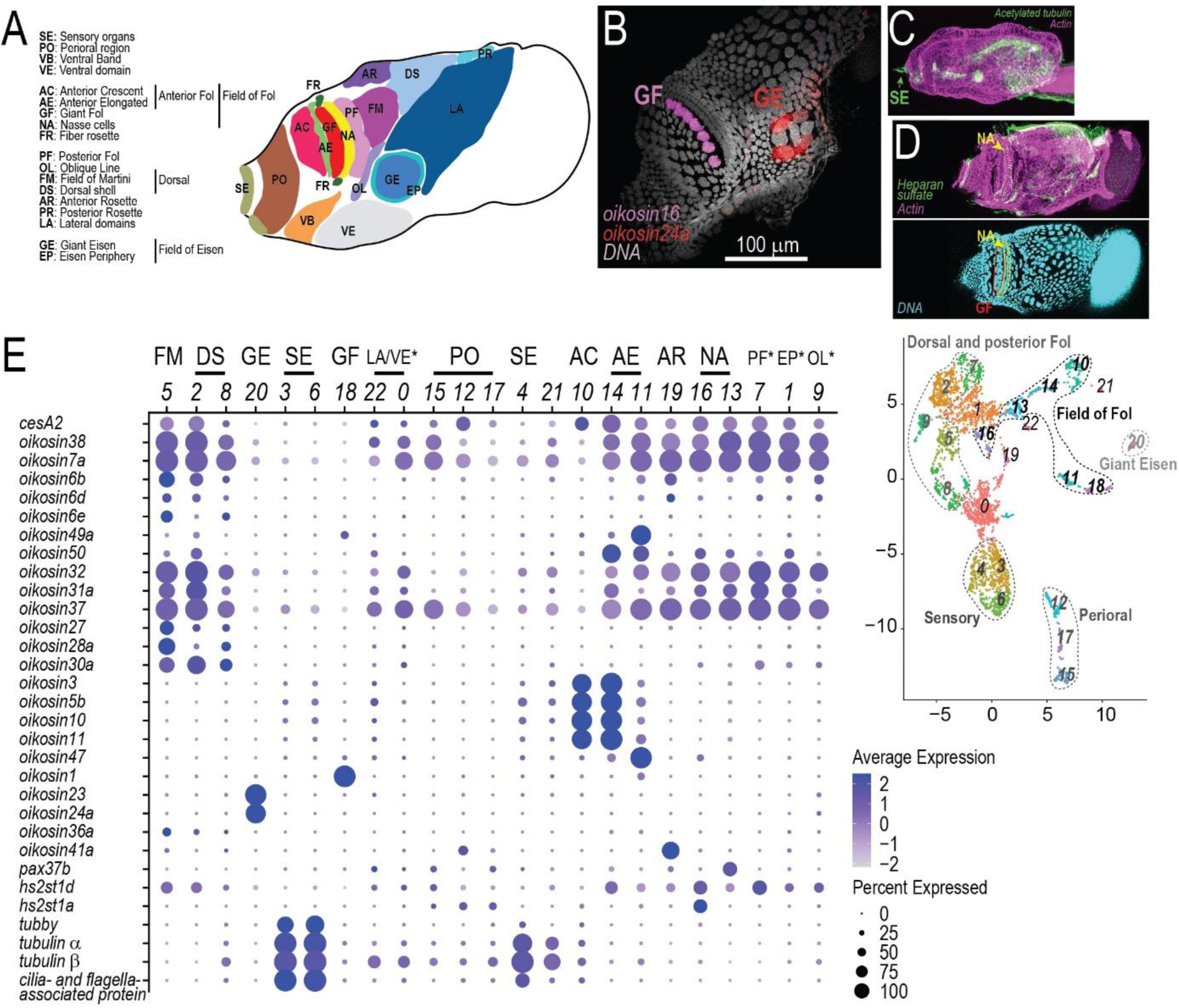
Diversity of oikoplastic cells in the mature epithelium. **A)** Representation of oikoplastic cell territories on a lateral view of the *O. dioica* adult trunk. **B)** Detection of oikosin gene expression in the *O. dioica* adult with HCR fluorescence, showing giant cells of the field of Fold and the field of Eisen. **C)** Immunostaining of acetylated tubulin in an *O. dioica* juvenile, showing cilia in the ventral sensory organ. **D)** Immunostaining of heparan sulfate in the *O. dioica* adult. The arrow indicate the anterior-most row of Nasse cells. **E)** Annotation of oikoplastic cell populations. The dotplot shows the gene expression in groups of cells produced by clustering *cesA2*-positive transcriptomes in *O. dioica* adult scRNA-seq libraries, which are represented in low-dimensional space in the right-side plot. Columns represent expression values for each group, abbreviations on top correspond to cell territories shown in **A)**. Genes were selected based on characteristic expression patterns marking cell territories of the mature epithelium [13, 27–29]. Stars indicate an annotation supported by extra markers shown in **FigS1**.

The field of Fol is a distinctive structure located in a dorsolateral position on the trunk, which includes different types of cells arranged in successive rows along the antero-posterior axis. We identified giant Fol cells (Fig 2A/E, “GF”) located in the central row with the expression of *oikosin14-21*, whereas the expression of *oikosin9-13* marked clusters corresponding to anterior Fol cells (Fig 2A/E, “AF”). We could also distinguish a cluster that expresses *oikosin47*, corresponding to elongated cells immediately adjacent to giant Fol cells (Fig 2A/E, “AE”). Nasse cells located posterior to giant Fol cells (Fig 2A/E, “NA”) could not be assigned using *oikosin* expression alone, as the diagnostic marker *oikosin48* was not found in the transcriptomes. We used other information for revealing two clusters representing the Nasse cells. The first one contains cells expressing *pax37B*, whose transcription in the epithelium is known to be restricted to Nasse cells [13] (Fig 2E, *cluster 13*). Cells present in the second one have high level expression of O-sulfotransferase transcripts (Fig 2E, *cluster 16*), consistent with high concentrations of heparan-sulfate fibrils detected with immunostaining in the anterior-most row of Nasse cells (Fig 2D). Dorsal territories include the anterior rosette, the posterior Fol and the oblique line cells, which we identified with the expression of *oikosin41*, *oikosin46* and *oikosin51*, respectively (Fig 2A/E, “AR”, “PF” and “OL”). Cells corresponding to the field of Martini were annotated based on higher expression levels of *oikosin2, oikosin27-28*, and *oikosin36a*, and they cluster together with cells expressing other oikosin transcripts that characterize the dorsal shell (Fig 2A/E, “FM” and “DS”).

The field of Eisen is another remarkable territory located in a ventrolateral position, consisting of seven giant cells surrounded by a ring of small cells. Giant Eisen cells were identified with the expression of *oikosins22-24* (Fig 2A/E, “GF”). Surrounding cells were identified with the expression of *oikosin2*, *oikosin7*, *oikosin25* and *oikosin50*, combined with reduced expression of dorsal and Fol-specific oikosins (Fig 2A/E and **FigS1**, “EP”). Similarly, lateroventral cells located posterior to the field of Eisen showed weak expression of *oikosin2* but strong expression of *oikosin37* (Fig 2A/E and **FigS1**, “LA/VE”). The expression of *oikosin2* is stronger in populations of cells corresponding to the perioral epithelium, which also expresses *oikosin37* but none of the oikosins that characterize the field of Eisen.

The expression of *cesA2* was highest in the anterior Fol and in the dorsal shell, and lowest in perioral and lateroventral cells. Giant cells showed weakest levels of *cesA2* transcripts, suggesting their main contribution to house synthesis could be structural proteins, and not necessarily cellulose. Cellulose fibrils are instead provided by adjacent cells such as Nasse cells with stronger *cesA2* expression, consistent with cellulose staining observed in the pre-house [28]. The lowest levels of *cesA2* and *oikosin* expression were measured in three cell populations, which are otherwise enriched for transcripts of the ciliary machinery. While they remain closer to the surface epithelium than to other cell types of the *O. dioica* adult, such expression pattern suggests these cells correspond to mechanoreceptors with prominent cilia that are found in sensory organs (Fig 2C).

We next compared how expressions of genes representative of diverse functions, including structural (glycosylation, extracellular matrix, glucan metabolism) and regulatory ones (transcription factors, signal transduction, cell cycle progression), vary among oikoplastic cells. For this analysis, we first identified transcripts that are preferentially expressed in the oikoplastic epithelium by comparing *cesA2*-positive cell clusters to other cell populations of the *O. dioica* adult (excluding all clusters represented by less than 500 cells) (Fig 3A/B).

**Figure 3:**
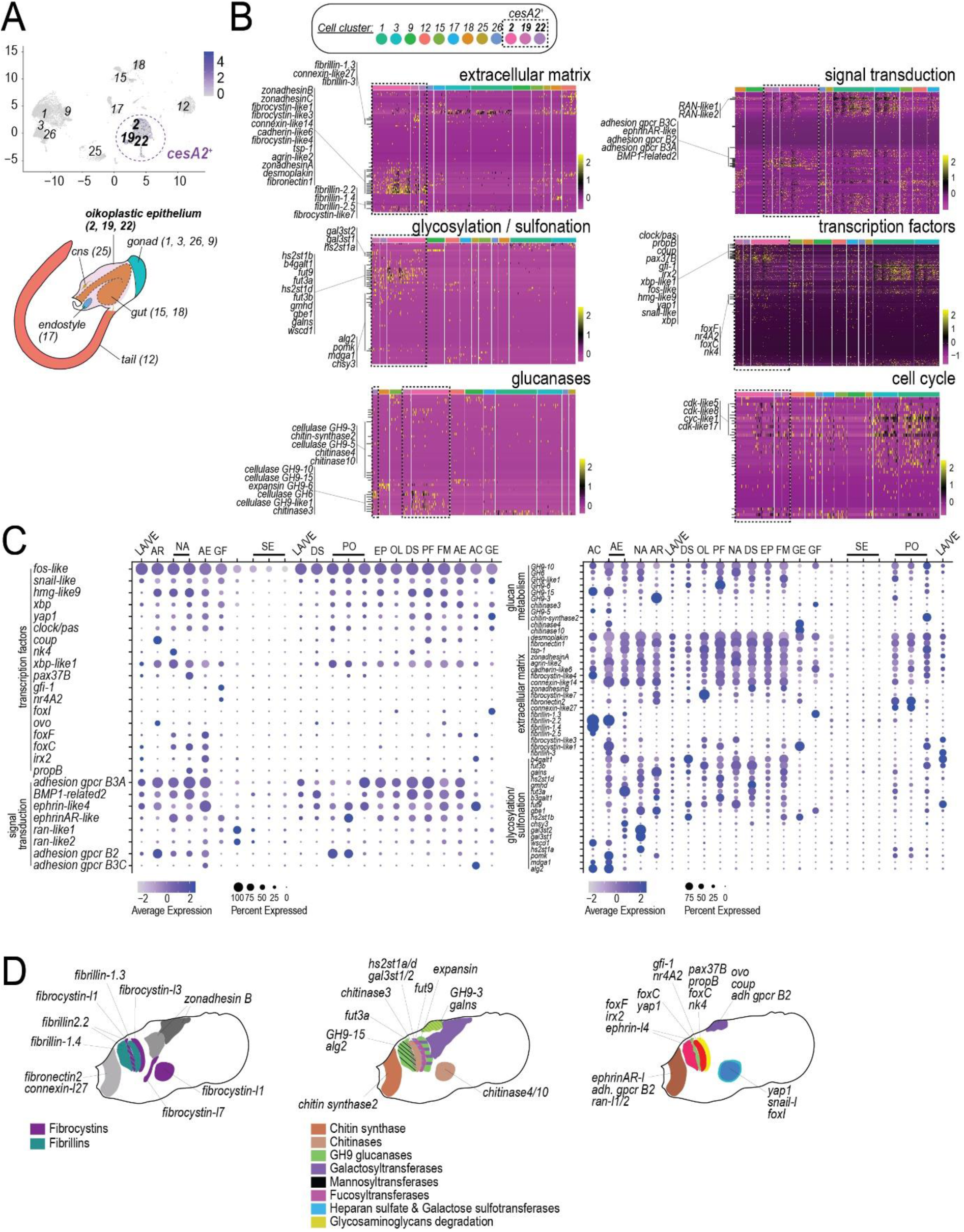
Functional patterning of oikoplastic territories. **A)** Top, representation of adult cell transcriptomes in low-dimensional space. Numbers indicate the cell clusters retained for differential gene expression analysis between *cesA2*-positive cells (bold) and other cell types. Bottom, localization of cell clusters in adult *O. dioica* tissues. **B)** Heatmaps showing the variation of gene expression between *cesA2*-positive cells and other cell types. For each functional category, labelled rows indicate top-expressed genes in *cesA2*-positive cells. **C)** Dotplots show the variation of gene expression among annotated oikoplastic cell types. **D)** Representation of gene expression patterns in oikoplastic cell territories on a lateral view of the trunk.

Out of 177 extracellular matrix (ECM) protein genes predicted in the genome, we found 25 preferentially expressed in the oikoplastic epithelium (Fig 3B, “extracellular matrix”). These include five homologues of fibrillin, which in vertebrate compose the elastic fibers together with elastin (not found in the *O. dioica* genome), and four fibrocystin genes out of the eight homologues found in *O. dioica*. We could not attribute a clear homology to the 14 remaining genes, but they all contained domains with predicted ECM functions. Sustained expression of these genes in oikoplastic cells may contribute to strengthening the oikoplastic ECM, which must endure mechanical constraints imposed by large differences in cell size and tension from pre-house secretion. Accordingly, transcripts encoding fibrillin and fibrocystin variants are largely restricted to fields of Fol and Eisen (Fig 3C, “AC”, “AE” and “GE” on right panel), territories containing cells with various sizes that are arranged in complex patterns [28]. These specific gene expressions constitute the first indications about the structural molecules that could be involved in shaping cell morphologies in the oikoplastic epithelium [11].

We found multiple genes predicted to encode enzymes that participate in heparan sulfate and glycosaminoglycans synthesis in the *O. dioica* genome (Fig 3B, “glycosylation/sulfonation”). It remains unclear whether these biopolymers contribute to either the ECM, the house, or both, but the systematic presence of this class of genes among the top-expressed ones in oikoplastic cells suggests critical functions. The dorsal shell, the elongated cells of the field of Fol and the Nasse cells show highest expression levels for glycosylating enzymes (Fig 3C, “DS”, “AE” and “NA” on right panel). Their expression is weakest in the anterior crescent of the field of Fol and in giant cells, which instead show strong expression of enzymes predicted to transfer sulfates on gangliosides (*wsdc1*) and heparan (*h2st1b*) enzymes (Fig 3C, “AC”, “GF” and “GE” on right panel). Using an immunoassay on whole-mount adult specimen, we detected a strong production of heparan sulfate localized in the first row of Nasse cells, demonstrating a functional specialization among cells whose morphology is otherwise undistinguishable (Fig 2D). The synthesis of cellulose necessarily involves the control of polymer length with specific glycoside hydrolases. Enzymes that cleave the β-1,4 glycosidic bonds in cellulose belong to the glycoside hydrolase family 9 (*GH9*), whose genes have expanded significantly in the *O. dioica* genome (supplemental information **SI2**). In *O. dioica*, the family includes a member with an expansin domain, unusual in animal GH9 but commonly found in bacterial cellulases involved in plant wall degradation [30]. The expression of such *gh9-expansin* is enriched in the posterior Fol cells, where it could play a function in relaxing cellulose fibers (Fig 3C, “PF” on right panel). We also observed that perioral cells and giant Eisen cells express high levels of chitin synthase genes (Fig 3C, “PO” and “GE” on right panel). Altogether, our results indicate a high level of specialization for producing different polysaccharides in the surface epithelium, that could contribute to distinct structures of the house (Fig 3D).

Most oikoplastic cell types share the expression of a set of genes with predicted regulatory functions during transcription (Fig 3B, “transcription factors”). It includes a *fos-like* gene and homologues of *snail*, *xbp*, *yap1*, *clock*, and one of the two genes with a single HMG domain present in the genome, *hmgl9*. On the other hand, we found genes whose expression is strongly restricted to one oikoplastic cell type, including *pax37B* in Nasse cells, *foxI* in giant Eisen cells, *gfi-1* and the nuclear receptor *nr4A2* in giant Fol cells (Fig 3C, “NA”, “GE”, “GF” on left panel). Transcripts of *foxC*, *foxF* and *irx2* are detected in multiple cell types, but they are preferentially expressed in territories of the field of Fol, reminiscent of their expression pattern during early development [14]. Out of four adhesion G-Protein Coupled Receptors (aGPCR) predicted in the genome, three are preferentially expressed in oikoplastic cells (Fig 3B, “signal transduction”). These proteins play important functions, notably in vertebrate CNS, both as cell adhesion molecules and components of signaling pathways [31]. The expression of *aGPCR-B3A* is shared among different cell types, but *aGPCR-B2* is preferentially expressed in the anterior rosette and in the perioral region, while the expression of *aGPCR-B3C* is restricted to anterior crescent cells of the field of Fol (Fig 3C, “AR”, “PO”, “AC” on left panel). Many of these genes are robustly transcribed in oikosin-expressing cell populations of the 8hpf juvenile, and in the next sections, we will examine how their expression correlates with the succession of developmental stages in larvae.

### Gene expression programs associated with early development

Cell lineage tracing showed that fate restriction occurs extremely early in *O. dioica*, with precursors of the main tissues of the larva such as the notochord, tail muscles and the nervous system identified after the fourth cleavage in the 16-cells embryo [32]. The earliest developmental stage examined in our study corresponds to embryos collected between the sixth and seventh cleavage, at 2hpf. Clustering the scRNA-seq dataset resolved transcriptionally distinct cell populations that we could identify as progenitors of ectodermal, mesodermal, endodermal and germ lineages. The proportions of cells processed here from different tissues and those described during cell lineage tracing are similar (Fig 4A and **B**). Except for *vas4*-expressing cells identified as the germline (Fig 4B/C), most clusters resolved at this stage show weaker expression of maternal transcripts and more zygotic ones (Fig 4D and **E**), suggesting that samples were collected soon after the first wave of zygotic gene activation [33].

**Figure 4:**
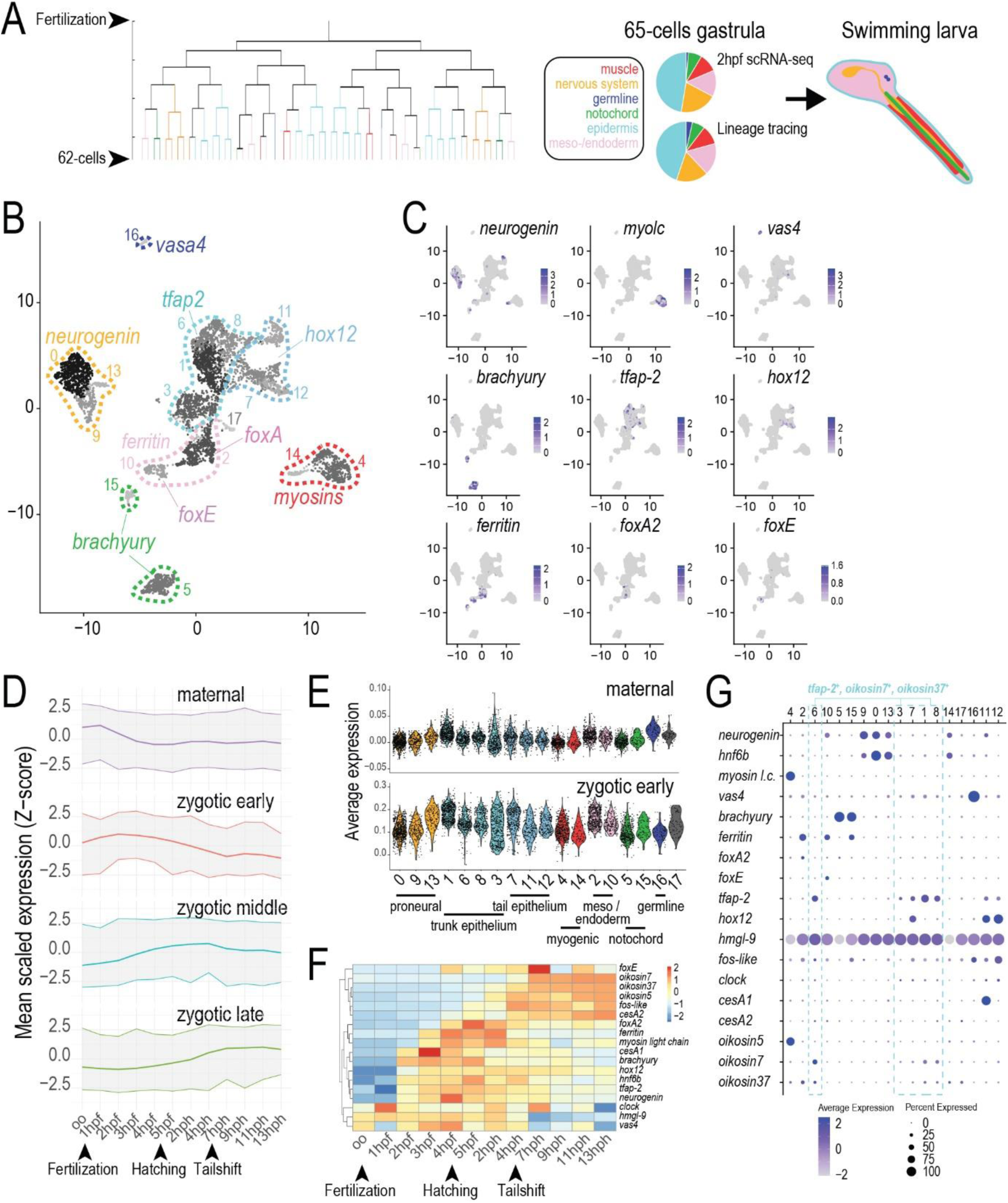
Transcriptomic signatures associated with early fate restriction. **A)** Left, fate map of *O. dioica* adapted from Stach *et al.* [32], with colors corresponding to different tissues identified in the swimming larvae. Right, proportion of cells progenitors for different tissues, measured in the annotated scRNA-seq library or in the 65-cells gastrulae of cell tracing experiments [32]. **B)** Representation of 2hpf single-cell transcriptomes in low-dimensional space. Progenitors corresponding to different tissues are indicated based on marker gene expression. The color code is the same as in **A)**, numbers in the chart area indicate cell clusters. **C)** Dotplot showing the expression of marker genes in 2hpf single-cell transcriptomes. **D)** Gene expression dynamics during *O. dioica* embryogenesis. Plots represent the mean expression of group of genes, clustered based on time series RNA-seq from bulk samples. The top plot shows genes strongly expressed in unfertilized oocytes, bottom plots show genes transcribed before 1h post-fertilization (zygotic early), between 3 and 4h post-fertilization (zygotic middle) and between 2 and 4h post-hatching (zygotic late). **E)** Expression of maternal and early zygotic genes in single-cell transcriptomes at 2hpf. **F)** Expression time course of gene marking different germ layers and the oikoplastic epithelium, based on bulk RNA-seq data. **G)** Dotplot showing the expression of gene markers in cell clusters annotated at 2hpf. Dashed lines indicate cell clusters corresponding to oikoplastic progenitors.

Using developmental genes conserved among chordates, we could attribute pro-neural and pro-myogenic identities to cells forming two highly distinct clusters in low-dimensional space (Fig 4B/C). Neural progenitors were identified based on strong expression of *neurogenin* and typical transcription factors involved in neural development such as *celf*, and *hnf6b*. In contrast, markers of neuronal differentiation like *elav* and *atonal* were not expressed. Muscle cells progenitors were identified by an expression of contractile machinery genes such as myosins and troponins.

The expression of the *brachyury* ortholog marks a highly distinct cluster of probable progenitors of the notochord (Fig 4B/C). Based on the ISH expression of a *ferritin* homolog, we can propose that a subset of *brachyury* positive cells are progenitors of the endodermal strand. Distinct endodermal lineages could be further pointed out with expressions of *foxA* and *foxE*, known to pattern the gut and the endostyle in larvae. These distinct transcriptome profiles show that at this early stage, meso-endodermal territories already include divergent lineages leading to the digestive tract. Similarly, we could identify among ectodermal cells the progenitors of different types of surface epithelium. Ectoderm progenitors are primarily defined with the expression of *tfap-2*, which in ascidians participates in larval epithelium development and cellulose synthase *cesA* regulation [34] (Fig 4B/C). In *O. dioica*, we observed that *tfap-2* remains co-expressed with *hmgl-9, oikosin7* and *oikosin37* throughout early development, suggesting that cells with strong *tfap-2* expression must include progenitors of the oikoplastic epithelium (Fig 4G and next section). The expression of *hmgl-9* appears also stronger in these candidate progenitors, possibly due to a contribution from maternal transcripts (Fig 4F). In contrast, transcripts of other markers such as *clock* and *tfap-2* derive primarily from early zygotic expression.

We identified *hox12*-positive cells as a second lineage of ectoderm progenitors (Fig 4B/C), that will give rise not only to the tail epithelium, but also to other structures located posterior to the trunk such as the Langerhans mechanosensory receptors [22]. This annotation is supported by the co-expression of several markers whose transcripts are highly restricted to posterior structures in larvae. These include the *cesA1* paralog of *O. dioica* cellulose synthase genes, whose expression has been observed only in epithelial cells of larvae [26]. We detected *oikosin5* transcripts in muscle progenitor cells, but the level of expression in this population decreases until 3.5hpf (Fig 4G, *cluster 4* and next section). The reason for this expression is unclear, since from 3.5hpf onward, *oikosin5* expression will remain restricted to cells that also co-express other markers of the mature epithelium. We observed a similar case with weak *oikosin37* expression in cell populations identified as *foxA2*-positive endodermal precursors (Fig 4G, *cluster 2*). We cannot exclude that these “ectopic expressions” could contribute to developmental functions.

### Molecular patterning of the tailbud embryo and differentiation of oikoplastic territories

Transcriptional profiling of scRNA-seq libraries at 2.5hpf and 3hpf suggests progressive reinforcement of cellular identities, while marks of terminal differentiation were observed in the 3.5hpf library only. The 3.5hpf dataset corresponds to the tailbud embryo, a developmental stage where morphological differentiation along the anteroposterior axis is first visible with major structures such as the rows of tail muscle cells and the notochord. Accordingly, lineages initially defined at 2hpf are now clearly resolved in low-dimensional space (Fig 5A). Neural progenitors form a compact and transcriptionally coherent domain characterized by the expression of markers such as *syt-II*, *GAD* and *VAChT*, indicating progression toward neuronal differentiation. This neural territory also includes subsets of cells expressing *pax6*, *hnf6b*, and *cdx3*, consistent with early patterning of the neural tube [35] and anteroposterior neural identities. Among mesodermal derivatives, we can now distinguish tail muscle cells - whose transcriptome is consistent with nascent myotome organization, and a smaller adjacent cluster expressing the *hand* cardiac marker. Transcripts detected *in situ* at 3.5hpf allowed us to define a distinct domain located at the boundary between the trunk and the tail. This domain includes epithelial cells with *cesA1* expression [26], consistent with the detection of cellulose fibrils protruding from the surface of tailbud embryos (Fig 5C). It also includes cells that express *pax2/5/8a*, which could correspond to precursors of cells associated with Langerhans receptors [36].

**Figure 5:**
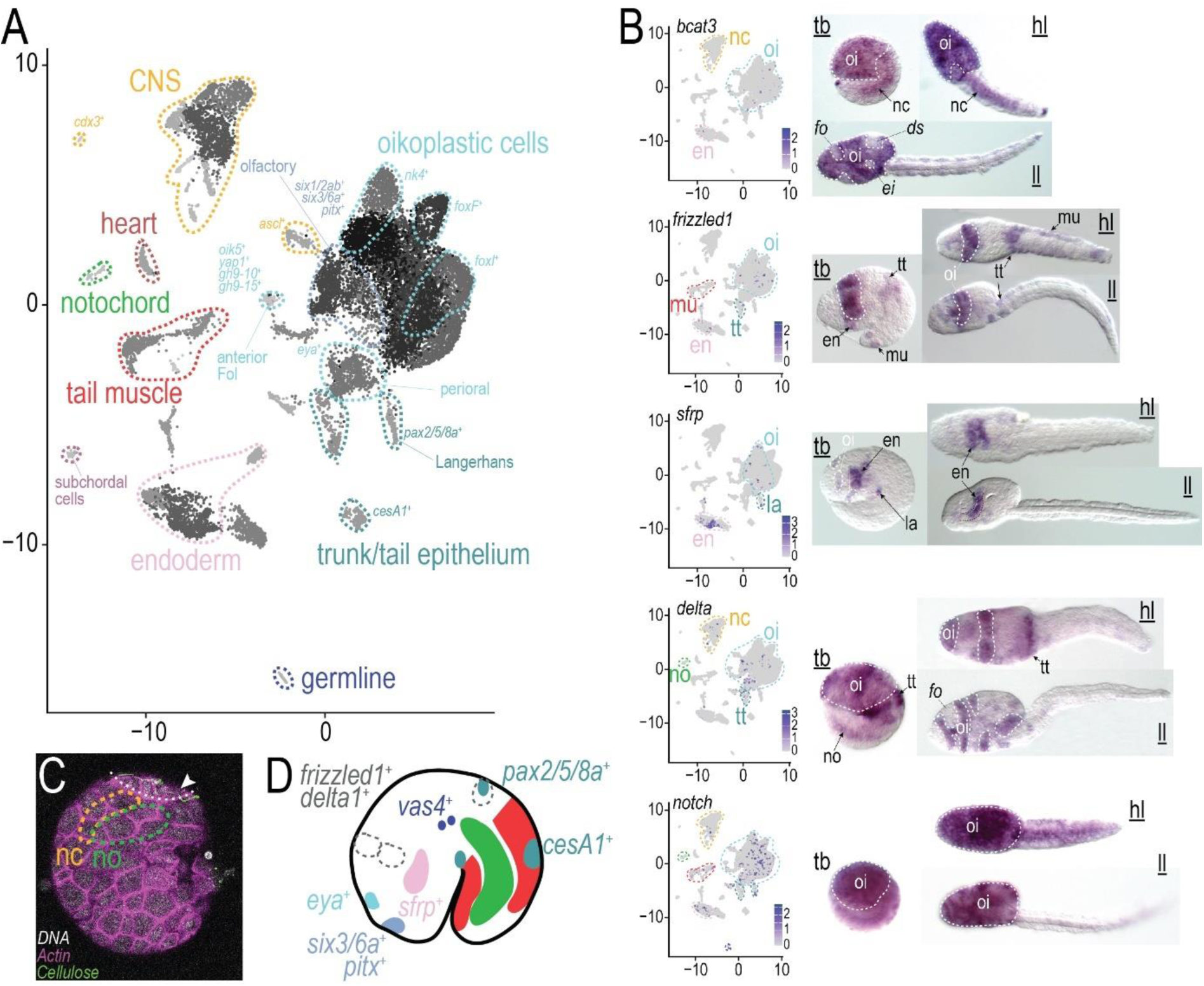
Molecular patterning during organogenesis. **A**) Representation of 3h30pf single-cell transcriptomes in low-dimensional space. Progenitors corresponding to different tissues are indicated based on marker gene expression. **B)** Expression patterns of signaling pathways genes. For each gene, we compare the average expression in 3h30pf single-cell transcriptomes (left panel) to *in situ* expression patterns (right panel) observed in either tailbud embryo (<u>tb</u>), hatched larva (<u>hl</u>) and larva just before the tailshift (<u>ll</u>). en, endoderm; ds, dorsal shell; ei, field of Eisen; fo, field of Fol; oi, oikoplastic progenitors; la, Langerhans receptor; mu, tail muscle; nc, nerve chord; no, notochord; tt, trunk/tail domain. **C)** Staining of a tailbud embryo showing cellulose fibrils (arrowhead) on tail ectoderm cells (marked with white dasheed line). Colored dasheed lines represent the position of the nerve chord and the notochord. **D)** Representation of gene expression domains in the tailbud embryo, including expression patters reported by Bassham et al. [36, 37]. Green and red areas mark the notochord and tail muscle, respectively.

The epithelium primordium of the tailbud embryos is patterned by homologues of Wnt signaling components, such as *bcat3*, *frizzled1*, and a secreted frizzled-related protein *sfrp* (Fig 5B). While their precise function in *O. dioica* remains unknown, expression at this stage is consistent with known functions of the Wnt pathway in chordate development, and we could detect transcripts in various tissues undergoing morphogenesis, such as the nerve chord and the anterior gut primordium. Except for *sfrp*, gene expressions also pattern the epithelium of the swimming larva, showing some overlaps with boundaries of oikoplastic cell territories.

Transcriptomes of the tailbud stage revealed a sharp increase in expression for several markers of the mature oikoplastic epithelium (Fig 6A, <u>3h30pf</u>). Transcripts of *fos-like*, *clock*, *tfap-2*, *hmgl-9* are enriched in populations of cells that include on one hand, oikoplastic progenitors marked with the expression of *oikosin37*, and on the other hand, progenitors of perioral olfactory organs where we detected the expression of *pitx*, *six1/2b* and *eya* [37] (Fig 5A and Fig 6A, *cluster 3* and *cluster 10* at <u>3h30pf</u>). The expression pattern of *fos-like* changed significantly between 3hpf, where transcripts are enriched in the tail ectoderm (Fig 6A, <u>3hpf</u>), and 3.5hpf where progenitors of the trunk epithelium represent most of cells positive for *fos-like* expression. Remarkably, *fos-like* expression becomes higher in tail tissues again at 4hpf, suggesting successive waves of transcription activation (Fig 6A, *cluster 12, 17, 22* and *14* at <u>4hpf</u>). For *clock* and *bmal* however, the transcription increase that starts at 3.5hpf remains restricted to progenitors of the trunk epithelium throughout larval development. From 3.5hpf, the expression of *tfap-2* gradually decreases in tail tissues and by 6hpf, it is restricted to oikoplastic cells which, at this stage, are engaged in the terminal steps of differentiation (Fig 6A, <u>6hpf</u>).

**Figure 6:**
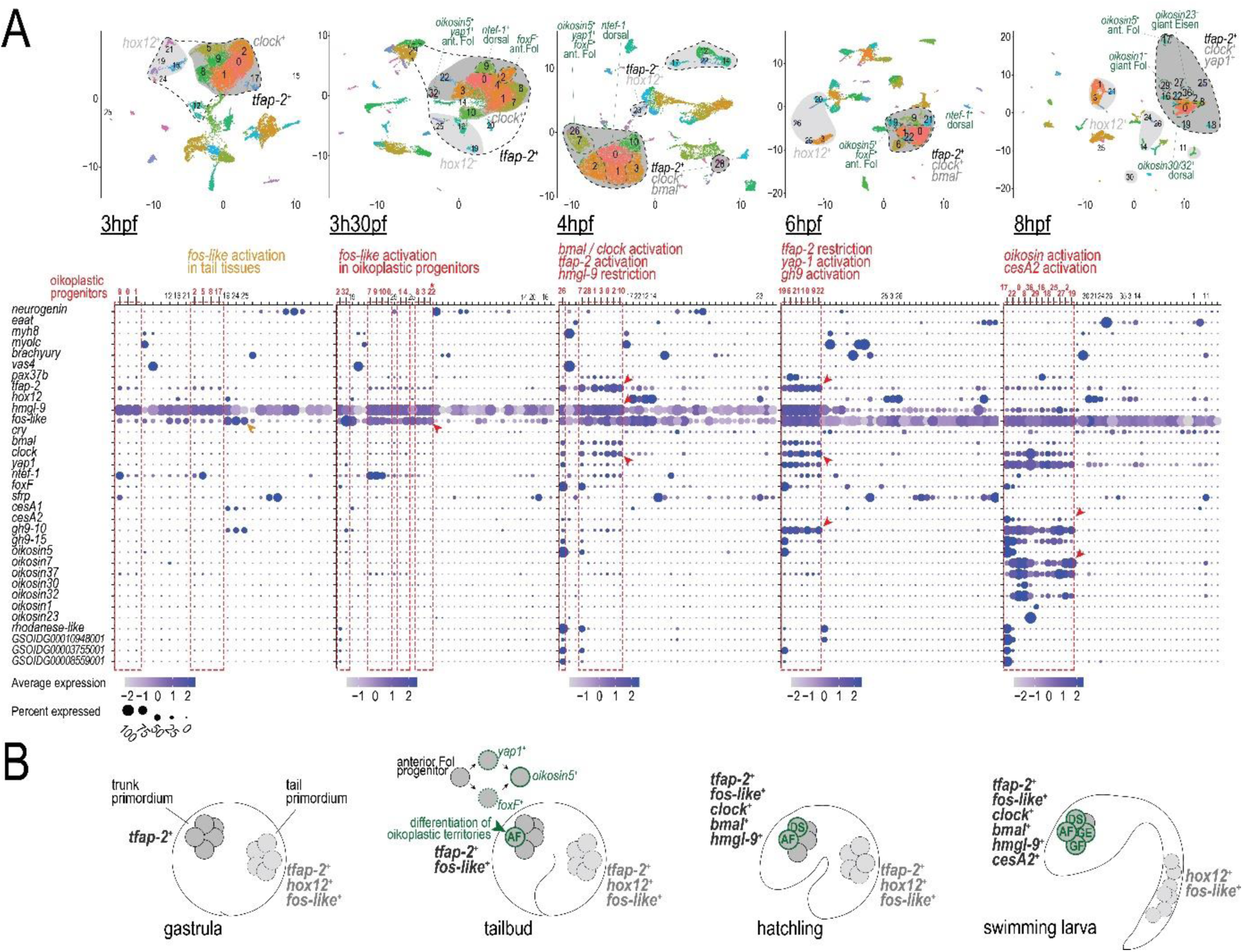
Transcription programs associated with the differentiation of the oikoplastic epithelium. **A)** Top panel shows single-cell transcriptomes from successive developmental stages, represented in low-dimensional spaces. Black dashed lines indicate epithelial progenitors positive for *tfap-2* expression. Dark and light grey areas indicate subpopulations of oikoplastic and tail epithelium progenitors, respectively. Dashed green lines indicate progenitors of distinct oikoplastic fields. Bottom panel show average expression of marker genes in cell populations represented in the top panel. Dashed red lines indicate oikoplastic cell progenitors. The star indicates a cell population that expresses both oikoplastic and neural marker genes. **B)** Representation of transcription programs active during epithelium development in the embryo.

Genes associated with oikoplastic ECM and house synthesis are largely inactive at this stage. However, in a small population spatially distinct from other progenitors, we could detect a gene expression pattern characteristic of the mature anterior Fol, which includes *oikosin5*, the transcription coactivator *yap1*, and the cellulases *GH9-10* and *GH9-15* (Fig 6A, *cluster 32* at <u>3h30pf</u>). Such early expression of *oikosin5* is consistent with ISH results showing it is the first oikosin gene activated during larval development [14]. Other markers that characterize the mature anterior Fol - generally orphan genes, were restricted to another population of *foxF*-positive cells included in the bulk of oikoplastic progenitors (Fig 6A, *cluster 2* at <u>3h30pf</u>). The separation disappears at a 4hpf embryos, where we can clearly reconcile expressions of *oikosin5* with other markers of the mature anterior Fol. We propose that *yap1*- and *foxF*-positive cells correspond to successive intermediates that could arise independently during the differentiation of anterior Fol cells (Fig 6B). We identified another population of tailbud stage progenitors with the expression of *ntef-1*, that will later mark dorsal oikoplastic cells (Fig 6A, *cluster 9* at <u>3h30pf</u> and *cluster 0* at <u>8hpf</u>). Even though molecular signatures of other oikoplastic territories remain elusive in the tailbud embryo, we cannot exclude that their progenitors may already be determined at this stage.

The variety of transcriptional signatures and their link with mature oikoplastic cells clearly indicate that oikoplastic progenitors in the tailbud embryos are already undergoing differentiation leading to terminal identities associated with anterior Fol and dorsal shell. This is taking place significantly earlier than morphogenetic processes responsible for organizing the surface epithelium, that have not been observed in animals younger than hatched larvae [14, 38]. The terminal differentiation of oikoplastic progenitors could involve the successive activation of *fos-like*, and *clock/bmal*, before the deployment of regulatory programmes that are specific for distinct cell territories [14] (Fig 6B).

In vertebrates, *clock* functions as an activator for the transcription of cryptochromes, blue light-sensitive flavoproteins that control circadian metabolic fluctuations. The *O. dioica* cryptochrome homologue *cry1* starts being expressed in diverse cell populations at 3.5hpf, but transcripts levels in oikoplastic cells remain systematically lower throughout development - except for giant cells. This pattern is remarkably similar for several other flavoproteins and for the FAD (flavin adenine dinucleotide)-producing enzyme riboflavin kinase (Fig 7A). Reducing the concentration of FAD destabilizes flavoproteins and can impact circadian gene function [39]. To test a role in the formation of the oikoplastic epithelium, we exposed *O. dioica* embryos to mild FAD concentration (50 μM) immediately after hatching and throughout larval development. In most cases, the treatment did not prevent the formation of a functional oikoplastic layer with distinctive cell territories (Fig 7B). However, more than half of treated animals exhibited a reduction of the size of the anterior Fol territory, invariably associated with a small deformation of the anterior trunk visible at the anal region. In a minority of animals, we could observe severe defects that often abolished the boundaries of cell territories usually visible in the oikoplastic epithelium. These strong phenotypes were remarkably restricted to the anterior trunk and did not impact the formation and maturation of posterior structures such as Langerhans receptors or tail tissues. These results show that perturbing the FAD balance during development produces specific phenotypes in the oikoplastic epithelium.

**Figure 7:**
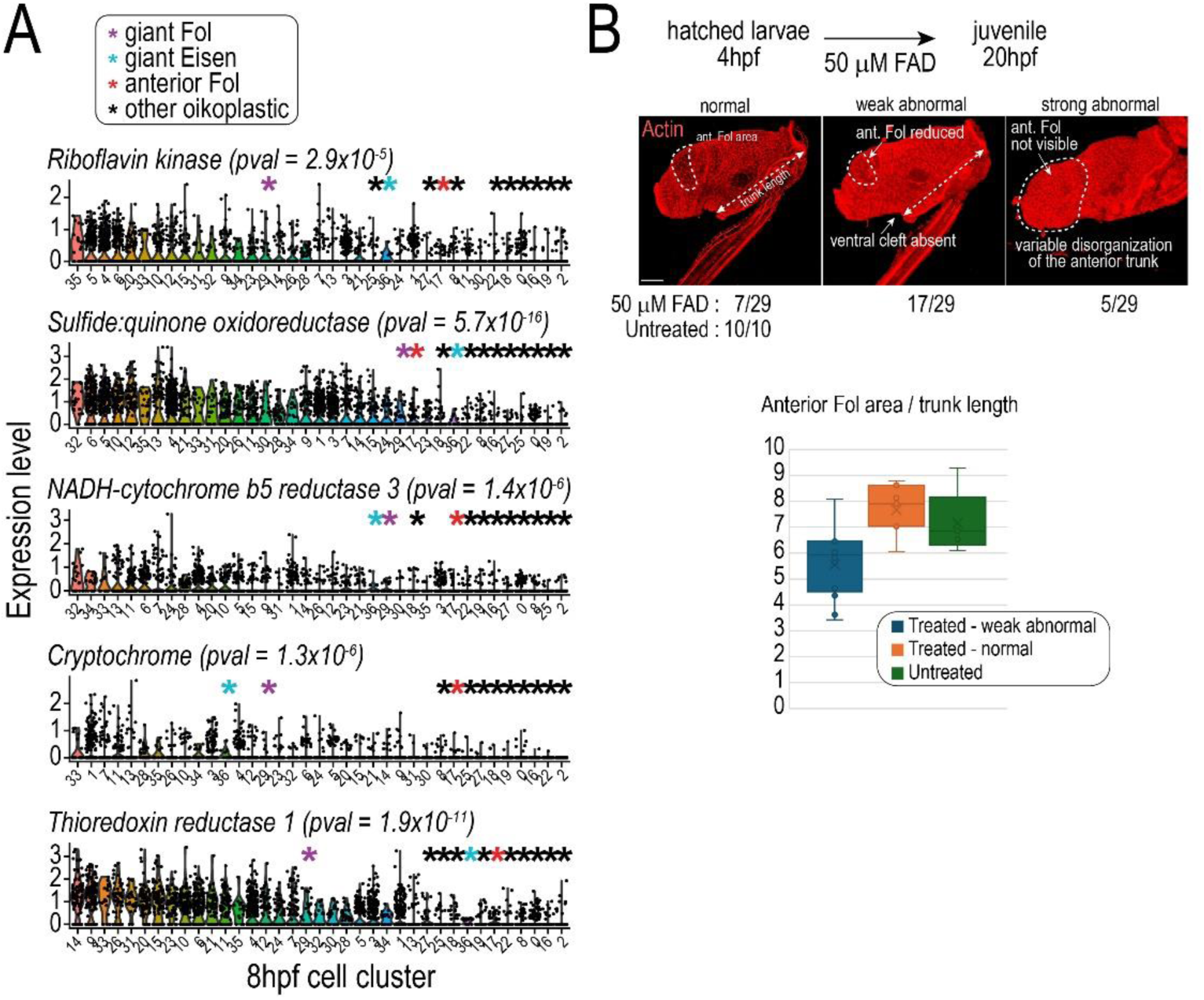
Flavoproteins play specific roles during the development of the oikoplastic epithelium. **A)** Comparison of the expression levels of flavoprotein genes between cell populations identified in the 8hpf scRNA-seq sample. Stars indicate clusters corresponding to oikoplastic cells. For each gene, we show the statistical significance of expression change between oikoplastic cells and all other clusters. **B)** Top, developmental defects caused by FAD exposure are apparent in a majority of samples. Weak morphological abnormalities consist in a reduced anterior Fol, that is systematically associated with the loss of the ventral cleft in the anal region. Strong abnormalities are represented by variable disorganization of the trunk epithelium, with one or multiple territories of oikoplastic cells being unrecognizable. Bottom, measurement of anterior Fol area relative to the trunk length, expressed as the distance from the anus to the gonad primordium.

### Acquisition of functionally specialized transcription programs during tailshift

After hatching, we can distinguish two well-defined phases before the completion of somatic development. The pre-tailshift phase (Fig 8A, steps (1-3)) corresponds to a conserved, successive formation of mature structures [40]. The notochord, tail musculature and tail innervation develop first after hatching, and swimming behavior usually begins shortly after leaving hatching. Next, ectoderm- and endoderm-derived tissues form primordia at diverse locations corresponding to oikoplastic cell territories [14, 38], the endostyle, the pharynx and the brain. The trunk anatomy is finally organized with the development of the hindgut and the posterior trunk, which involves cell migration from the tail [24]. During this succession of events, the mouth develops progressively and it opens when foregut development is completed [41]. The tailshift phase corresponds to a repositioning of the tail orientation relative to the trunk, to achieve the conformation observed in adult animals (Fig 8A, step (4)). Sometimes referred to as a metamorphosis [42], this event is critical for tail functions related to house inflation and filter-feeding. Mechanistic and molecular information remains scarce, however, and we decided to take advantage of our developmental series to examine the transcriptomic changes associated with the tailshift transition.

**Figure 8:**
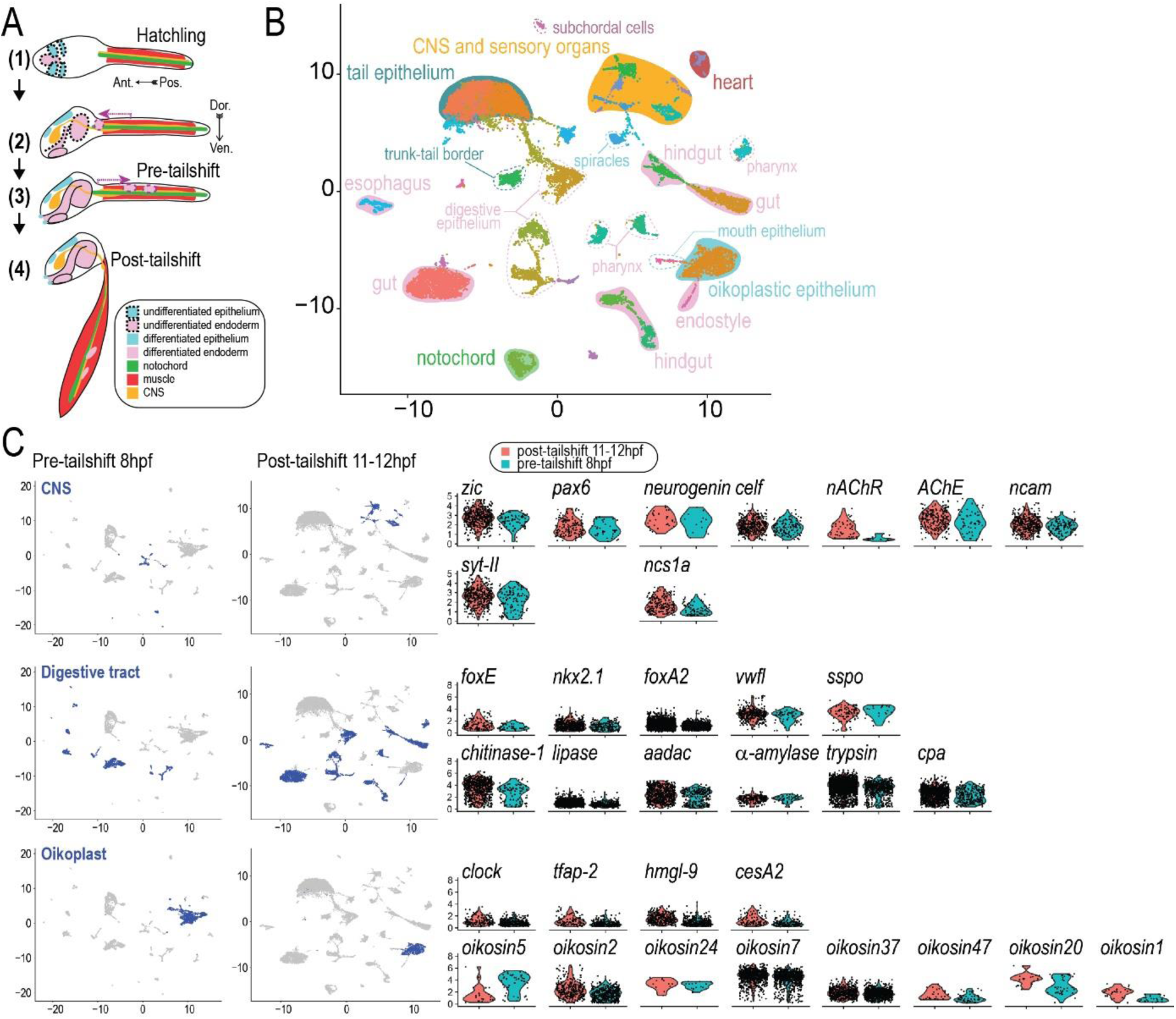
The tailshift transition. **A)** Schematic representation of morphogenetic processes and organs in the newly hatched larva (1), during early (2) and late (3) organogenesis, and in post-tailshift juvenile (4). Purple arrowheads in (2) and (3) indicate migrating cells that respectively contribute to gut development and subchordal cells. **B)** Representation of 11-12hpf post-tailshift transcriptomes in low-dimensional space showing annotated cell populations. **C)** Gene expression changes associated with the tailshift transition. The panel on the left shows cell populations selected for comparison of gene expressions between pre- and post-tailshift samples. The panel on the right shows gene expression levels between selected cells from the CNS, the digestive tract, and the oikoplastic epithelium.

Samples collected at 8hpf in our dataset have undergone all morphogenetic processes associated with the pre-tailshift phase. This is evidenced by reduced expression of *ferritin*, a strong marker of the endodermal strand that is not expressed in the mature hindgut. Clustering reveals many distinct cell populations with marker genes corresponding to well-specified structures such as oikoplastic territories, the notochord, and specialized cells of the endostyle, the gut, and the nervous system (Fig 8C). At this stage, however, differentiation markers such as *neurogenin*, *pax6,* and *zic* in the nervous system, *foxA* in endodermal tissues, *nkx2.1* and *foxE* in the endostyle remain strongly expressed.

Samples collected at 12hpf exhibited the distinctive morphology of post-tailshift juveniles. The transcriptomic landscape shifts markedly toward functional maturation, with the most prominent changes observed in cells of the digestive tract. At this stage, cells that correspond to distinct segments of the gut segregate into clearly resolved identities (Fig 8B). Cells allocated to the same organ, for example the pharynx and the hindgut, get further separated into two distinct clusters. The composition of top-expressed genes reflects the emergence of cell types with specialized secretory functions related to digestion. Compared to pre-tailshift transcriptomes, expressions of several hydrolase genes were upregulated (Fig 8C, right panel), including trypsins, chitinases, metallopeptidases, carboxypeptidases and amylases, confirming further that the tailshift transition is tightly connected with the acquisition of digestive functions. This is also reflected in the transcriptomes of endostyle cells, where strong expression of sco-spondin, *von Willebrandft factor-like* gene, and lectin genes suggests mucus secretion contributing to process food particles. Patterns of *oikosin* genes expression that characterize the mature epithelium are now fully deployed, consistent with the onset of house production. Genes associated with synaptic function (*syt-II*), neurotransmitter synthesis (*AChE*), and neuronal signaling (n*AChR*) are strongly expressed in cell populations annotated as neural, consistent with the establishment of neuronal circuitry required for coordinated tail movement and sensory processing.

## Discussion

The small size of larvaceans precluding most dissection techniques to isolate individual organs, scRNA-seq represents an exclusive approach for revealing cell diversity in these organisms. By gaining such resource, we could bring exhaustive transcriptomic information for *O. dioica* tissues, whereas past knowledge was based mainly on selected candidate genes. The fact that we could link most scRNA-seq transcriptomes to an existing organ, either in developing animals or in adults, supports a successful capture of cell diversity. Female and male gamete represent an exception, and their absence in the dataset could be a consequence of unusual cell size and/or fragility. We are confident that our dataset will be pivotal for future localization of new cell types in the *O. dioica* anatomy, in combination with other tools such as sensitive probes and high-resolution imaging [43]. This is illustrated with the oikoplastic epithelium, whose various cell groups characterized at the histological level are clearly reflected among single-cell transcriptomes. The segregation of these populations into distinct sub-groups, as shown for example with Nasse cells, strongly suggests multiple cell types with specialized functions could often be present within the same oikoplastic territory. How larvacean houses trap particles, and how these structures are processed in the marine environment, have emerged as key issues for understanding the roles of larvaceans in the ecosystem [44–46]. We could reveal the expression of many genes that participate in the metabolism of diverse polysaccharides in oikoplastic cells, suggesting that other critical components than cellulose may be present in the larvacean house. Although we cannot exclude other roles in organizing the oikoplastic cell layer, such chemical diversity could contribute to prey recognition or pathogen neutralization inside the house during filter-feeding.

The scRNA-seq resource will be instrumental for raising comparisons with other taxa for which single-cell transcriptomes are currently available. The representation of adult samples is a central asset here, which is highly relevant for uncovering key cell types and molecular function associated with life history strategies. In a developmental perspective, the sampling at short time intervals from pre-gastrulation stages until hatching will be useful to address the conservation of transcription programmes involved in patterning the chordate body plan. Our work constitutes a decisive advance for this objective by providing scRNA-seq data for developmental stages in close succession until hatching. With the examples shown in this study, we illustrate how our dataset may allow to trace every cell lineage of *O. dioica* at the molecular level, from the blastula stage until the completion of somatic development. Deciphering the regulatory logic of developmental programmes may require complementary experiments to monitor chromatin regulation in *O. dioica*, such as chromatin conformation capture and sc-ATACseq. Here, we chose to narrow our analyses on molecular processes associated with key features of larvacean biology, which include on one hand the development of the house-producing organ and on the other hand, rapid organogenesis during larval stages. Indications gained only with gene expression patterns limit the possibility to derive strong mechanistic knowledge, but nevertheless, they allowed us to enrich our understanding of larvacean biology by a large extent.

Previous research showed, with candidate approaches, that lineage-specific homeodomain genes duplicates often show specific expression in the larval oikoplastic epithelium [8], and they are required for the proper organization of cell fields [13, 14]. For most duplicates, such expression remains transient and does not persist in the mature epithelium for most lineage-specific duplicates. We observed the same dynamic expression for *tfap-2*, with a significant decrease in oikoplastic cells after larvae have undergone tailshift. In fact, the expression of *tfap-2* is also well correlated with the developmental progression of the tail epithelium, since transcripts levels in posterior tissues already decrease between 4 and 6hpf, a time window where tail morphogenesis becomes complete. These are strong indications that in larvaceans too, *tfap-2* contributes to the development of ectoderm-derived tissues [34] which include the oikoplastic eptihelium. However, our analyses indicate that the differentiation of ectodermal progenitors into either tail- or oikoplastic epithelium requires the superimposition of specific transcription programmes. Consistent with results gained in ascidians [47], our results suggest that *hox12* should be required for the differentiation of tail ectoderm in *O. dioica*. For the differentiation of the oikoplastic layer, we suggest new roles for *hmgl-9* and circadian genes, that add to the functions characterized for homeodomain genes [13, 14]. Deciphering the transcriptional cascades that connect *hmgl-9*, *tfap-2*, *clock* and *hox12* will represent a significant step to elucidate how oikoplastic cells could diverge from an ancestral epithelium, and to test further their proposed relationship with neuroepithelial identities [13]. With this example in mind, we firmly believe that together with complementary approaches, examining other tissues represented in our dataset will also prove instrumental for investigating the origins of core processes that characterize chordate development.

## Supporting information

SI1:List of genes mentioned in the manuscript

SI2:glycoside hydrolases of the Oikopleura dioica genome

## Acknowledgments

We thank Marios Chatzigeorgiou for the critical reading of our manuscript. This study was supported by a grant for the Sars Centre core budget (NFR grant 234817 “Sars International Centre for Marine Molecular Biology Research, 2013–2022”) and a grant to Daniel Chourrout. (NFR grant 250005 “accelerated evolution in chordates and the origin of larvaceans”), both from the Research Council of Norway.

